# Loom response in mouse superior colliculus depends on sensorimotor context

**DOI:** 10.1101/2024.12.06.627189

**Authors:** Stefano Zucca, Auguste Schulz, Pedro J. Gonçalves, Jakob H. Macke, Aman B. Saleem, Samuel G. Solomon

## Abstract

Visual motion is produced both by an organism’s movement through the world, and by objects moving in the world such as potential predators. Choosing appropriate behaviour therefore requires organisms to distinguish these sources of visual motion. Here we asked how mice integrate self-movement with looming visual motion by combining virtual reality and neural recordings from superior colliculus (SC), a brain area important in visually-guided approach and avoidance behaviours. We first measured locomotion behaviour and neural activity while animals approached an object in virtual reality, and while the same object loomed at them. In both cases, vision dominated activity in superficial layers (SCs), while locomotion had more influence on activity in intermediate layers (SCim). In addition, animals instinctively slowed their locomotion when nearing the object, or when the object neared them. To directly test animals’ ability to distinguish self-from object motion we replayed the visual images generated during object approach. Locomotion behaviour often changed during replay, showing animals are able to establish if visual motion is matched to their self-movement. Further, decoders trained on locomotion behaviour, or on population activity in SC, particularly in SCim, were able to reliably discriminate epochs of replay and object approach. We conclude that both mouse behaviour and SC activity encode whether looming visual motion arises from self-or object movement, with implications for understanding sensorimotor coordination in dynamic environments.

**Highlights:**

- We recorded from superficial (SCs) and intermediate (SCim) superior colliculus in VR
- Vision dominated SCs, while SCim was modulated by both vision and locomotion
- Mice altered behaviour when visual experience did not match that expected from their locomotion
- Population activity differed between matched and unmatched visual experiences, particularly in SCim

## Introduction

The meaning of sensory activity is often ambiguous, because multiple contexts produce identical patterns of sensory stimulation. For example, identical sequences of retinal images can be produced by an object moving (looming) towards an observer, or an observer’s movement toward that object. Accounting for this ambiguity requires organisms to establish whether a pattern of sensory stimulation matches that expected from self-movement, or is more likely to arise from object movement, such as a potential prey or predator. The importance of distinguishing sensory stimulation arising from self-or object movement to organism survival may explain why there are widespread neural correlates of self-movement, such as locomotion, that converge with other sensory signals in multiple brain areas ^1-5^.

Superior colliculus (SC) is one such multimodal brain area, that is implicated in visually-guided approach of potential prey or other objects, and avoidance of potential visual threats ^6,7^. SC has access to visual sensory signals directly from retina ^8^ and via visual sensory cortices ^9^, and can access locomotion-related and high-level contextual signals from multiple brain regions ^10,11^. Retinal and visual cortical inputs are biased to superficial layers of SC (‘SCs’). Non-visual inputs are instead biased towards intermediate (‘SCim’) and deeper layers, which are also more tightly linked to behavioural outputs (see ^7,12-15^ for reviews).

The anatomical connections suggest that SCs should be primarily sensitive to visual signals, while activity in SCim should be more dependent on the interaction of locomotion and vision, and more able to distinguish self-from object movement. There is, however, limited understanding of how locomotion and visual signals combine in SC. Visual responses in SCs are strong, but can increase or decrease during locomotion ^16-20^. Visual responses in SCim are weaker ^18,21-25^, and we know little about locomotion’s impact on SCim activity ^17,18^. In addition, these measurements have rarely been made in contexts where animal locomotion is relevant to the visual stimulus. To establish how SC responds during self- and object movement, we therefore measured activity in SCs and SCim in mice immersed in a virtual reality environment.

## Results

Mice were head-restrained over a treadmill, which we configured to control movement in a virtual reality (VR) environment. Mice could move along a 100 cm long platform in VR by moving on the treadmill (**Figs 1A,B**). We used 32-channel multielectrode arrays to measure spiking activity from populations of single-units in SCs (30 sessions in 7 mice) and SCim (35 sessions in 9 mice) (**Fig 1C**).

**Figure 1:**
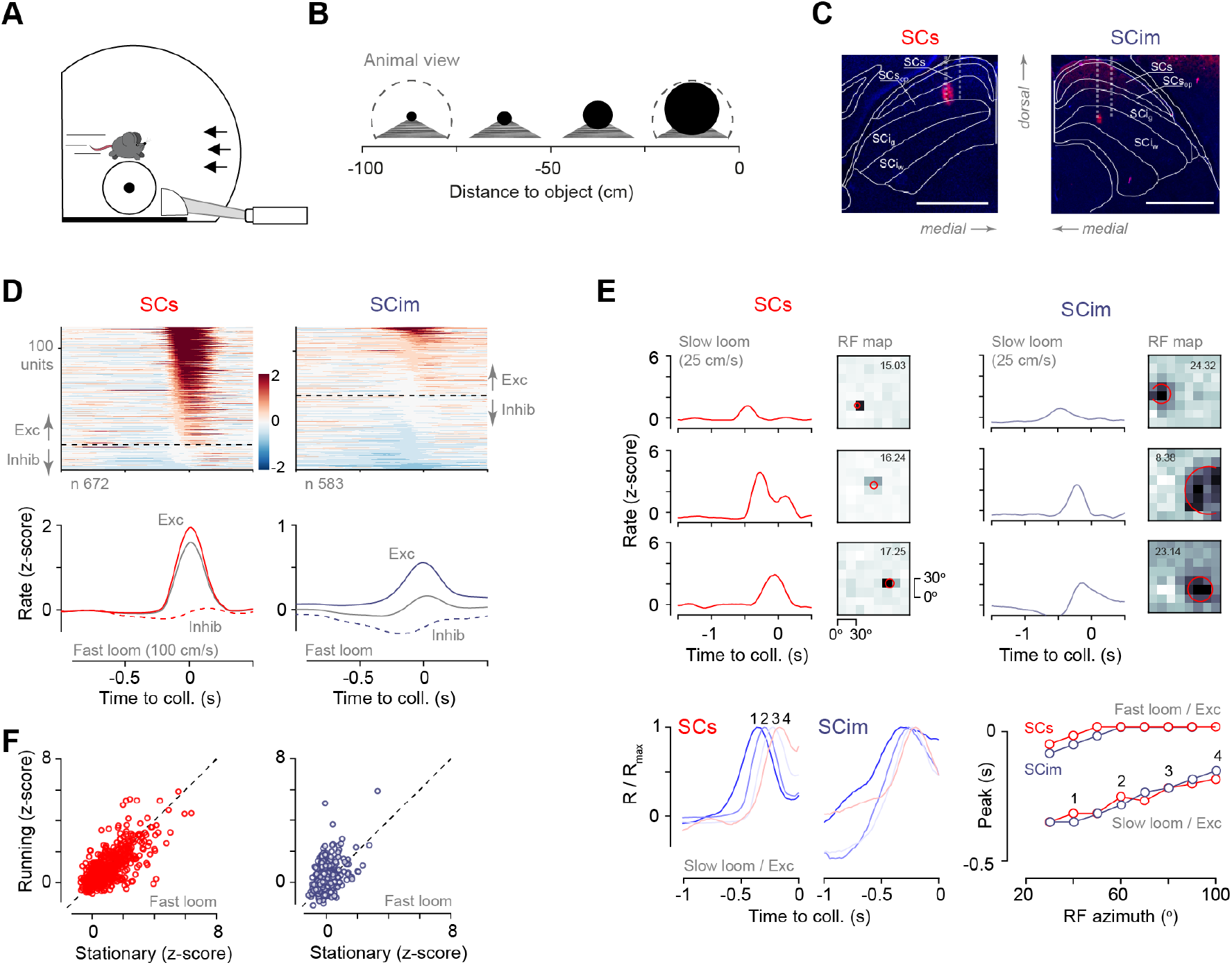
Loom response in SC combines visual and self-motion signals A. Animals were head-restrained above a treadmill in an immersive virtual reality environment. **B**. A round black object loomed towards the animal at constant speed. Schematics show the approximate view of the virtual object to the animal at each distance; dashed line indicates the visual limit of the virtual environment. **C**. Extracellular recordings were made from SCs and SCim using high-density probes. Images of sections through SC showing red fluorescence deposited by a DiI-coated electrode. Inferred electrode tracks are indicated by vertical lines. SCs image is flipped on the horizontal axis. **D**. Visual response is strongest in SCs. (*upper*) Responses (z-scores) of consistent units to object loom (100 cm/s), sorted by response in 0.2 s preceding collision. (*lower*) Average response over all units in SCs or SCim, and for units where looms increased (‘Exc’) or decreased (‘Inhib’, dashed line) activity. **E**. Visual receptive fields predict timing of loom response. (*upper*) Example units showing response to slow object loom (25 cm/s), and receptive field maps for black squares. Receptive field maps are flipped on the horizontal axis so that the nasal visual field is on the left, and temporal visual field is on the right. Darker indicates larger positive weight at that location. Red circle indicates estimates of location and size of excitatory receptive fields in each case. (*lower left)* Average responses to slow loom, for units where the looming object increased activity. Units were grouped by the azimuthal locations of their receptive field; the four lines of varying colour show average response in each of four of the azimuthal groups. (*lower right*) Time to response peak as a function of receptive field azimuth. Units with more nasal receptive fields (azimuth closer to 0) respond earlier to the looming object. Numerals indicate relevant curves in the lower left plot, from which the indices were derived. **F**. Locomotion has a more consistent impact on SCim. Response to fast object looms over the 0.2 s preceding collision, in SCs and SCim, during stationary and running conditions.

### Neurons in SCs and SCim respond to looming visual objects

We first measured response to an object that loomed at the animal. On each trial, a dark round object (8 cm diameter) appeared at the distant end of the platform and then moved towards the animal at a speed of 100 cm/s. We analysed responses where activity was consistent across trials (see Methods), yielding 672 units in SCs, and 583 in SCim (**Fig 1D**). We found that the looming object increased activity in most SCs units. Responses in SCim were more diverse, such that the looming object increased activity in some SCim units, but decreased activity in others. Consequently, activity was on average higher in SCs (mean z–score 0.85, s.d. 1.03) than in SCim (0.07, s.d. 0.56; *p* << 0.001, Student’s t-test).

Vigorous response to object loom in SCs is consistent with previous measurements that show highly responsive, spatially-localised visual receptive fields in most SCs neurons (e.g. ^18,26-29^). Conversely, fewer neurons in SCim show measurable visual receptive fields ^18,21,23^, potentially explaining reduced response to looming objects in SCim. To understand the relationship between loom response and receptive fields, we determined receptive field maps by presenting flashed black and white squares at different positions in the visual field, and fit the maps for each unit with two-dimensional Gaussians. In these recordings we could extract receptive fields for at least one luminance polarity in most SCs units (607/711, 85%; see Methods), and fewer SCim units (230/830, 28%) (examples in **Fig 1E**). Most of these units responded to the black squares, and we found that receptive field size (s.d. of the Gaussian) for black squares was smaller in SCs (mean 7.3 degrees, s.d. 3.9, n = 569) than SCim (mean 14.4, s.d. 6.2, n = 219; *p* < 0.001, Student’s t-test).

Visual neurons should respond when the object’s edge passes through their receptive field, and receptive field location should therefore predict the relative timing of neural responses to the loom stimulus. We tested this prediction in units with a receptive field map, which also increased activity in response to the looming object. We averaged loom response among units with receptive fields at similar positions along the nasal-temporal axis, and found the time at which loom response peaked (**Fig 1E**). Since objects looming at 100 cm/s expanded very rapidly, there was little difference in time to response peak across the visual field. We therefore also presented objects that approached at slower speed (25 cm/s; see also **Suppl. Fig 1**). At slower speeds, we found that loom response peaked earlier among neurons with more nasal receptive fields, in both SCs and SCim. The temporal pattern of responses to a looming object is therefore consistent with that expected from neurons’ receptive field locations.

### Self-movement modulates response to looming visual objects

Locomotion influenced the response of SC neurons to object movement. Animals moved sporadically during presentation of object looms, and so we defined each trial as either ‘running’ (locomotion speed at least 2 cm/s) or ‘stationary’ (less than 2 cm/s) and calculated average activity in each condition, over the 0.2 s preceding object collision. We found that locomotion could either increase or decrease activity in SCs neurons, with no net effect (**Fig 1F;** mean difference in z-scores of -0.05, s.d. 0.80; *p* = 0.141, paired Student’s t-test). In contrast, locomotion consistently increased activity in SCim, by a mean 0.23 (s.d. 0.72; *p* < 0.001; SCs vs. SCim: *p* < 0.001, Student’s t-test). Our measurements therefore reveal that individual neurons throughout SC process both visual and locomotion signals: visual response is strongest in SCs, but locomotion more consistently increases SCim response.

### Vision and self-motion signals also converge during object approach

Visual looming also arises when an animal approaches an object. To characterise response to looms caused by self-movement we made a simple change to the virtual reality (VR) environment. On each trial the same dark object still appeared at the distant end of the platform, but now it stayed there. As animals moved along the platform, their self-movement brought them closer to the object; the upshot was that the object again loomed in the visual field, but now that looming was caused by animal self-movement. Time to reach the object varied between trials, so here we analysed neuronal activity as a function of distance between animal and object.

We found a consistent pattern of response in SCs as animals approached the object in VR: population neuronal activity increased to a peak, and then subsided (**Figs 2A,B**). Object approach produced a different pattern of responses in SCim. Some neurons showed a distinct peak in activity as the animal neared the object. Other neurons showed distinct reduction in activity at similar positions. In most SCim neurons, however, we saw a ramp in activity (either increase or decrease) as animals moved along the platform. Activity patterns in these neurons usually showed a peak or trough as the animal neared the object. Overall, we saw a consistent pattern of response in SCs and a diversity of responses in SCim.

**Figure 2:**
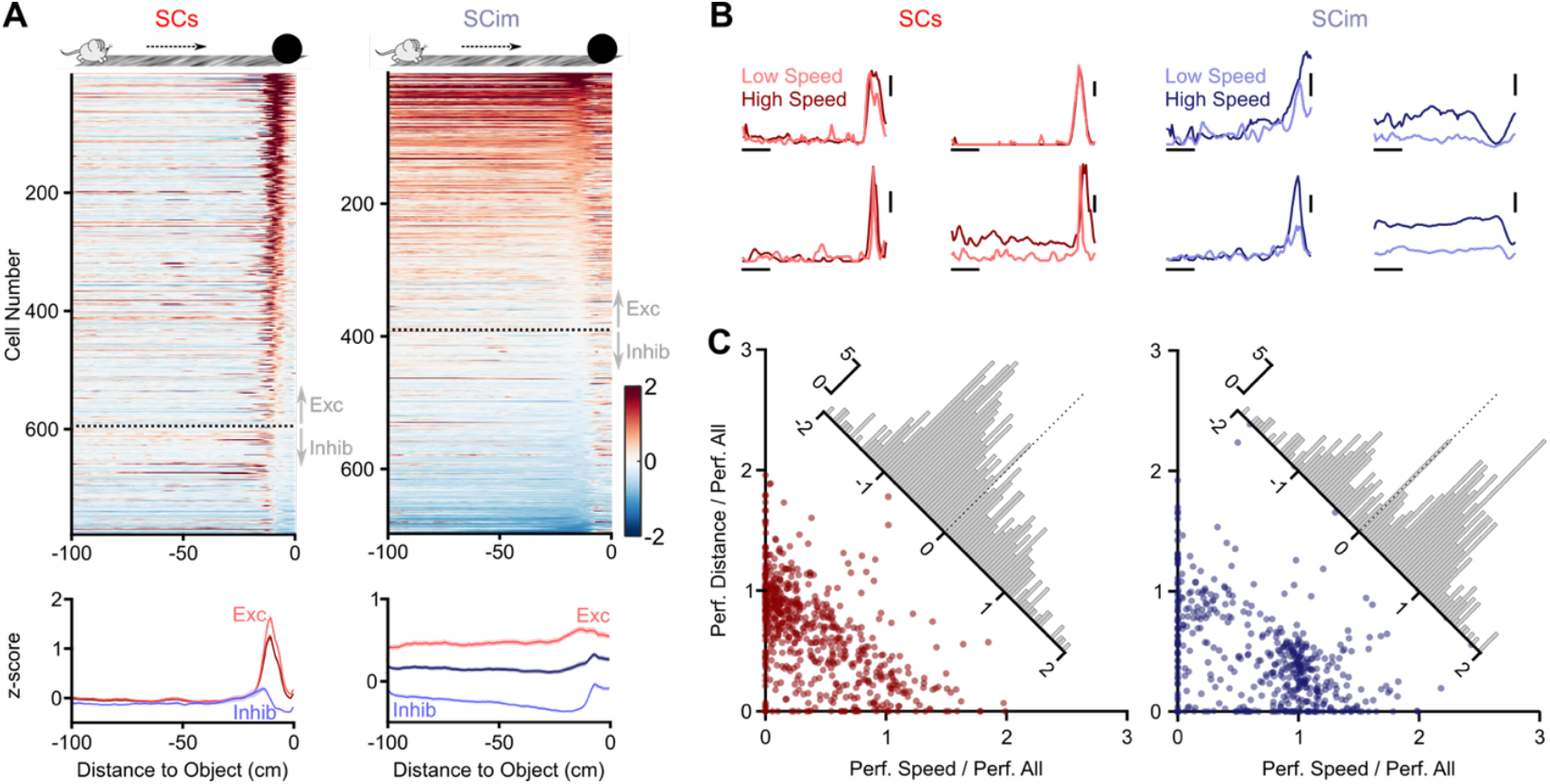
SCs is dominated by vision, and SCim is dominated by self-motion, during object approach. A. Patterns of activity were stereotypical in SCs, but diverse in SCim, as animals approached a stationary object at the end of the virtual platform. (*upper*) Response (z-scores) of consistent units, sorted by response in the 0–10 cm (SCs) or 10–20 cm (SCim) preceding collision. Colourbar indicates unit response in z- scores. (*lower*) Average response over all units in SCs or SCim, or over units in which object approach increased (‘Exc’) or decreased (‘Inhib’) activity. **B**. Speed of approach had more impact on activity in SCim and less impact in SCs. Example units showing average activity during trials of high locomotion speed (darker colours; 5 fastest trials), or low locomotion speed (lighter colours; 5 slowest trials). Horizontal scale bar 20 cm; Vertical scale bar 1 z-score. **C**. Relative contribution of locomotion speed and distance-to-object to activity in individual units in SCs (*left*) and SCim (*right*). Each point represents the performance (‘Perf.’; explained variance) of a ridge regression model when only locomotion speed (abscissa) or distance (ordinate) was allowed to vary, normalised to the performance of the model when both were allowed to vary. These normalized values could be greater than one due to the cross-validation. Histograms show the difference between these performance indices.

Locomotion speed influenced activity during object approach, particularly in SCim. SCim activity depended on approach speed throughout the trial, even early in the trial when the object was still distant (examples in **Fig 2B**). SCs activity, by contrast, was similar during both slower- and faster approaches. We used ridge regression to establish the relative contribution of visual and self-movement signals to SC activity during object approach. With these regressions we estimated the amount of variance in neural activity that could be explained by the distance between animal and object (equivalently, the location of object edges in the visual field), or by locomotion speed (see Methods). To assess the relative contribution of each factor, we normalized the cross-validated explained variance attained when using either locomotion speed or distance, to that attained when using both. Note that these normalized values can be greater than one due to the cross-validation. We found a variety of responses across the population, such that some neurons were more dependent on distance to object, while others were more dependent on locomotion speed (**Fig 2C**). Neurons in SCs showed stronger dependence on distance (median normalized explained variance = 0.706) compared to approach speed (0.256; *p* << 0.001, n = 627, Wilcoxon Rank Sum Test). Neurons in SCim instead showed stronger dependence on approach speed (median = 0.888) compared to distance (0.402; *p* << 0.001, n = 468).

### Vision drives instinctive locomotion behaviours

Mice were free to choose how they moved along the virtual platform. Locomotion behaviour was, however, dependent on vision and stereotypical: mice consistently slowed down as they approached the object (**Fig 3A**). This slow-down behaviour could be observed on the first day of exposure to an object, and then became more pronounced as animals gained experience and ran faster down the platform (**Fig 3C**). Slow-down behaviour was not limited to black objects - in separate experiments we found that animals showed the slow-down behaviour when they instead encountered a white object (not shown), and continued to show slow-down behaviour when the white object was subsequently replaced with a black one.

**Figure 3:**
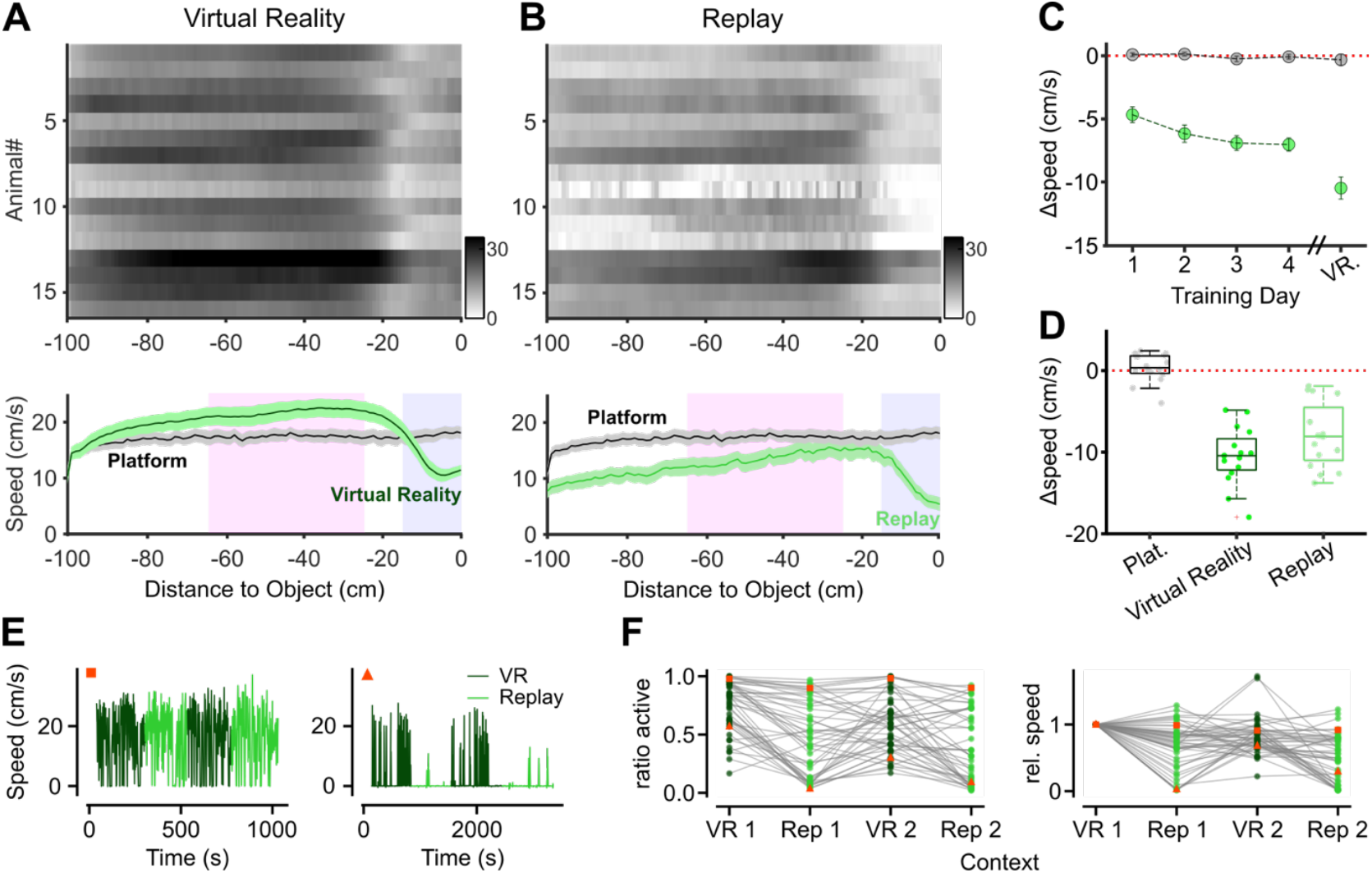
Vision and sensorimotor context drive instinctive locomotion behaviours A. (*top*) Animals reduce locomotion speed (slow-down) as they approach an object in virtual reality (VR). Each row of the image represents the locomotion speed of one animal, averaged across all VR trials in all post-training sessions. Colorbar indicates animal speed (cm/s). (*bottom*) Slow-down behaviour was present during object-approach but not when the object was absent (platform trials). Mean and s.e.m. of locomotion speed across animals. Shaded regions indicate locations used for estimates of locomotion speed used in panels C-D **B**. Same as A but during replay, where the visual stimulus was not predicted by current locomotion behaviour. Slow-down behaviour remained but overall locomotion speed was reduced during replay of object-approach trials. **C**. Slow-down behaviour emerged in the earliest stages of exposure to virtual objects. Each point shows locomotion speed (mean and s.e.m. across animals) when animals were near the object (blue-grey regions in panels A,B), relative to that at larger distances (pink regions in A,B). Equivalent measurements were made during platform trials. The last point shows the data from the post-training sessions shown in A. **D**. Comparison of slow-down behaviour in platform, VR and replay trials post-training. Small symbols show individual animals. **E**. Temporal dynamics of locomotion behaviour in two example sessions. (*left*) Session in which locomotion speed was similar during VR (dark green) and replay (light green) blocks of trials. (*right*) Session in which the animal ceased locomotion upon exposure to replay trials, and resumed locomotion when reintroduced into VR. **F**. (*left*) Fraction of time in each condition where animal locomotion speed was at least 0.1 cm/s. Data represent 48 sessions in 14 animals. Red squares and triangles indicate example sessions in E. (*right*) Average locomotion speed in each condition, relative to that in the first exposure to VR.

How does locomotion behaviour differ when visual stimuli match or conflict with that predicted by self-movement? To address this, we stored the sequence of visual images generated as an animal approached the object, and replayed those sequences later in the session. This allowed us to compare two conditions: locomotion behaviour during VR, where the visual stimulus matched that predicted by self-movement, and during replay, where it did not. We found that behaviour changed during the replay condition, such that animals often ran slower or even ceased moving during one or both replay epochs (**Figs 3E,F**, Wilcoxon signed-rank test; *p* < 0.02 for all conditions relative to the previous condition, n=48; see also **Suppl. Figs S2,S3**). When animals ran during the replay condition, their patterns of movement resembled that in the VR condition - animals slowed down when the object was close (**Figs 3B,D**). Similarly, presentation of independently looming objects (as in **Fig 1**) could evoke slow-down behaviour at each of the loom speeds we tested (**Suppl. Fig S4**).

The similar slow-downs observed for VR and replay conditions, and looming objects, suggests that slow-downs are an instinctive sensorimotor response to a looming visual stimulus (at least in head-restrained animals), whether that looming arises from self-movement or object movement. The changes in locomotion behaviour between VR and replay conditions, however, suggests that mice become aware - at some level - of whether the visual experience is a result of their own movement or not.

### SC activity is sensitive to match between vision and self-movement

We hypothesised that patterns of neural activity in SC, particularly SCim, depend on whether the visual stimulus is matched to self-movement. Because the temporal pattern of visual stimulation when presenting the looming object as in **Fig 1** (where object movement was smooth) could be very different to that during object approach (where object movement depended on animal speed), we tested the hypothesis by analysing behaviour and neural responses during the VR and replay conditions described above. Consistent with the overall impact of locomotion shown in **Fig 2**, SCim activity often reduced dramatically when the animal simply stopped moving during replay of the visual stimulus (**Suppl. Fig S5**). To provide a stricter test of the hypothesis, we restricted the analyses to sessions in which animals showed a similar range of locomotion in the VR and replay conditions (**Suppl. Fig S2**), and only when the animal was within 50 cm of the object. We divided each trial into non-overlapping 0.2 s epochs, and performed logistic regression to classify each measurement epoch as arising from VR or replay conditions, using behavioural and neural variables (**Fig 4A**, see Methods).

**Figure 4:**
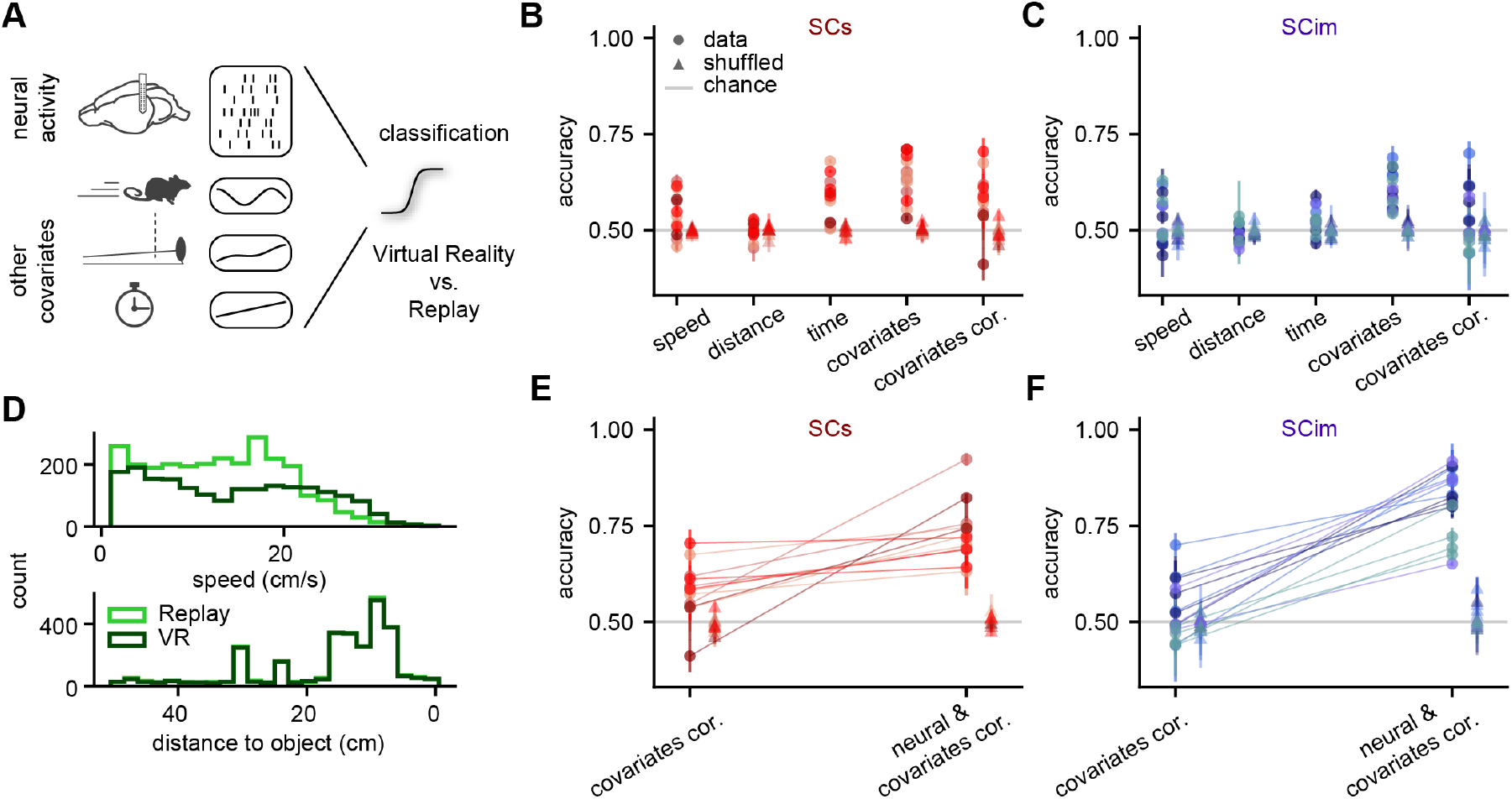
Neural activity in SC discriminates virtual reality and replay conditions A. Logistic regression was used to classify epochs of time as belonging to Virtual Reality (VR) or replay given neural activity from SCs or SCim and/or other covariates, including running speed, distance to object, or time into the experiment. **B**. Test classification accuracy of SCs sessions for individual covariates (speed, distance, time) and all covariates combined. ‘Covariates cor.’ indicates classification performance when using a corrected subset of the data (confining analyses to samples of VR and replay conditions with the same speed range, the same average time into the experiment, and confining analyses to distances within 50 cm of the object). Circles indicate mean performance in each session, and triangles indicate mean of shuffled controls for each session. Mean and s.d. Lines were obtained across four train-test splits. Symbols of the same hue are sessions from the same animal. **C**. Same as B but for SCim sessions. **D**. (*top*) Distributions of locomotion speed differs for VR and replay conditions. (*bottom*) Distributions of distance-to-object are matched, by design, in VR and replay. **E**. Test classification accuracy of SCs sessions when including all corrected covariates, or when including neural activity as well as those covariates. Symbols of the same hue are sessions from the same animal. **F**. Same as E but for SCim sessions.

We first measured classification performance when the regression model was blind to neural activity. Even in sessions where overall locomotion was similar, logistic regression was able to predict if the animal was in a VR or replay condition from combinations of behavioural variables, including locomotion speed, distance to the object, and time in the session (Wilcoxon signed-rank test; covariates: *p* < 0.001, n = 12 SCs sessions; *p* < 0.001, n = 15 SCim sessions; **Figs 4B,C**). Classification with locomotion speed alone was sometimes sufficient (speed: SCs sessions, *p* = 0.034; SCim sessions, *p* = 0.277). Time into the experiment also had some predictive power (time: SCs sessions, *p* < 0.001; SCim sessions, *p* = 0.035), but this may be an indirect effect of locomotion speed, as animals usually ran faster earlier in the session. Above chance classification persisted in many sessions even after ensuring average time into the experiment and locomotion speed ranges were matched between VR and replay conditions (covariates cor.: SCs sessions, *p* = 0.009; SCim sessions, *p* = 0.389). We tested the classifier after shuffling the trials between VR and replay conditions, and found that performance dropped to chance levels for all variables tested (covariates cor.: SCs sessions, *p* = 0.204; SCim sessions, *p* = 0.470). The distributions of locomotion speed in the two conditions remained different even after speed-matching (**Fig 4D**), so in the following we compare classification performance when including neural activity, to that obtained from the covariates alone.

We tested whether inclusion of neural activity during these measurement epochs would improve model performance (**Figs 4E,F**). We recorded spiking activity of populations of neurons alongside behaviour, and calculated the mean firing rate in the same epochs. We trained the classifier on subsets of the data for which measurement time and locomotion speed ranges were similar across VR and replay conditions. We found that neural activity in SCim improved classifier performance (mean ± s.d.: 56.1 ± 23.0 % increase, *p* < 0.001, n = 15 sessions; **Fig 4F**). Neural activity in SCs also improved classifier performance (28.8 ± 27.7% increase, *p* < 0.001; n = 12 sessions; **Fig 4E**), but to a lesser extent (SCim vs. SCs; *p* = 0.005, Mann-Whitney U test: 32.0). These improvements in SCim and SCs were not dependent on the range of distance-to-object included in the analyses (**Suppl. Fig S6**). Thus, neural activity in SC, particularly SCim, depends on whether the visual stimulus is matched to that expected from the animal’s locomotion.

## Discussion

We found that neuronal activity in SCs was dominated by vision, but that vision and self-movement signals contribute to neuronal activity in SCim, for both looming objects and during object approach. Animals instinctively slowed-down when an object was near, and further altered their behaviour when visual experience did not match that expected from their locomotion. This difference between visual experience and expectation from locomotion behaviour was also encoded by population neural activity in SC, particularly in SCim.

Mice often ceased moving during replay of the sequence of images they had previously generated while approaching an object. The behaviour likely reflects their instinctive ability to predict and account for the visual stimulation brought about by their self-movement. Virtual reality allowed us to generate a mismatch between predicted and actual visual stimulus, and mice were sensitive to this discrepancy. Neural activity in SC was also sensitive to this mismatch. Although our analyses revealed behavioural differences between VR and replay conditions, SC activity could still distinguish between the two conditions when behavioural differences were accounted for. We infer that both animals and neural activity in SC, particularly SCim, are able to tell the difference between visual stimuli produced by object- and self-movement, even when they give rise to identical patterns of visual stimulation. Previous work has described subpopulations of neurons in primary visual cortex that are sensitive to brief disruptions (mismatch events) between predicted and actual visual stimulus (^30^; but see ^31^). Similarly, response of neurons in visual cortical areas POR and LI (but not V1) to a small moving spot depends on whether the spot’s movement is coupled to animal locomotion ^16^. These latter cortical responses appear to depend on a subpopulation of SCs neurons (‘widefield cells’), and the pooled calcium signal of populations of widefield SCs neurons is increased when a small spot’s movement is uncoupled (either slower or faster) from animal movement ^16^. Widefield cells, which project from SCs to the pulvinar (lateral posterior nucleus), show weak responses to large visual stimuli like the looming object we presented ^28,32^, but we cannot rule out a contribution from widefield cells to the observed differences in VR and replay conditions.

Where comparison can be made our neurophysiological measurements appear consistent with previous work. Receptive fields were large in SCim units where we could recover them, as in previous work ^18,22-25^. SCim receptive fields were larger than those of most SCs neurons, but similar in size to widefield and ‘horizontal’ SCs cell classes ^26,28,32^. We also found weaker responses to visual looms in SCim than SCs. Response to visual looms is often measured after centering the stimulus on a neuron’s receptive field (e.g. ^23,33,34^), but here objects always loomed from in front of the animal, and the edge of the stimulus therefore always moved across the visual field in the nasal-temporal direction. Direction-selectivity is a common feature of receptive fields in SCs ^35^, but can also be found in deeper SC ^21^, so we cannot rule out the possibility that neurons in SCim (or SCs) would have been more responsive if we had tested looms centred at other locations.

Finally, the substantial impact of locomotion on activity in SCim seems consistent with very limited previous work (^18^; but see ^17^). We also found that locomotion had limited impact on net populational activity in SCs, consistent with previous measurements ^17-20^, though we note that specific subpopulations in SCs may show more consistent facilitation ^36^.

The relatively strong dependence of SCim activity on locomotion and related features is consistent with the differential inputs to SC layers. SCs receives strong visual input from retina ^8^, primary visual cortex and some secondary visual areas (LI, LM, P, POR; ^9^). By contrast, SCim receives weak if any retinal input and cortical visual inputs are mainly derived from other secondary visual areas (A, AM, PM, RL; ^9^) that may form a ‘dorsal’ stream through mouse visual cortex. SCim also receives multi-modal sensory input from auditory and somatosensory pathways (e.g. ^22,37^), as well as multiple non-sensory signals from subcortical and cortical areas (see ^10,11^) including inputs from retrosplenial cortex, prefrontal cortex and striatum among others (e.g. ^38-40^). These inputs to the SCim make it an ideal site for establishing the VR context, and broadcasting those signals.

In freely-moving mice, overhead looming objects usually induce escape when a refuge is present, and freezing behaviour when a refuge is absent ^41-43^. Looming objects in the frontal visual field can also induce escape to refuge, though usually after initial freezing behaviour ^44^. Our behavioural measurements show that head-restrained mice slow-down when presented with looming objects in the frontal visual field, whether or not that visual loom is matched to their locomotion. Other visual behaviours that are readily elicited in untrained, head-restrained animals include ‘vidgetting’ and locomotion changes produced by novel grating patterns ^45,46^; ‘locomotion arrest’ produced by flashed lights (^47^; see also ^48^); and ‘burrow ingress’ produced by overhead looms ^49^. Slow-down behaviour adds to this repertoire of instinctive behaviours in head-restrained animals; whether there is a relationship between them needs to be established.

Animals explored the virtual environment at moderate locomotion speeds of about 20 cm/s. The instinctive slow-downs we observed when animals were near an object may be adaptive, providing both time and motor state for a greater range of subsequent behaviours. Objects and agents (conspecifics, or potential threat or prey) approaching at slow speed will evoke a wave of activity across SC during this period (**Fig 1E**), analysis of which could allow animals to choose among appropriate approach or avoidance behaviours (cf. ^38^). Striking predators will likely move more quickly (e.g. ^50^), and a rapidly approaching immediate threat will near simultaneously activate a large fraction of visually responsive neurons in both SCs and SCim (**Fig 1D**). The different spatiotemporal dynamics of population activity during slower object approach and fast object loom may be sufficient to allow rapid avoidance behaviours in presence of immediate threat, while reducing the likelihood of false-alarms.

Our measurements leveraged immersive virtual reality to afford tight experimental control, while retaining expression of instinctive behaviours. Our findings demonstrate that both animals and neurons in SC can distinguish whether current visual experience matches that expected from the animal’s actions. The convergence of visual and locomotion signals onto SC, and the fact that SC is well placed to instruct behaviour, indicate that contextual variation in SC activity could be important in allowing animals to shape their behaviour.

## Resource availability

### Lead contact

Further information and requests for resources and reagents should be directed to and will be fulfilled by the Lead Contact, Samuel Solomon.

### Materials availability

This study did not generate new unique reagents. Commercially available reagents are indicated in the ‘Key resources table’.

## Acknowledgements

This work was supported by the Biotechnology and Biological Sciences Research Council (BBSRC; BB/R004765/1), by the UKRI Frontier Research Grant (EU underwrite; EP/Y024656/1) and by the Human Frontier Science Program (RGY0076/2018). This work was also supported by the German Research Foundation (DFG) through Germany’s Excellence Strategy (EXC-Number 2064/1, PN 390727645), the German Federal Ministry of Education and Research (Tübingen AI Center, FKZ: 01IS18039). AS is a member of the International Max Planck Research School for Intelligent Systems (IMPRS-IS). We thank Sarah Ruediger for comments on the manuscript.

## Author contributions

This work was conceptualized by S.Z, A.B.S. and S.G.S.; experiments were performed by S.Z.; formal analysis and visualization by S.Z., A.S. and S.G.S.; formal logistic regression analyses and visualisation by A.S., P.J.D. and J.H.M.; original draft writing by S.Z, A.S., A.B.S. and S.G.S.; review and feedback by all authors; supervision and funding acquisition by S.G.S, A.B.S and J.H.M.

## Methods

### Experimental model and subject details

All experiments were performed in accordance with the Animals (Scientific Procedures) Act 1986 (United Kingdom) and Home Office (United Kingdom) approved project and personal licences. Mice (n = 16 C57BL6/J male wild-type, age 12-16 weeks) were housed in groups of maximum five under a 12-hour light/dark cycle, with free access to food and water. All electrophysiological recordings were carried out during the dark phase of the cycle.

### Method details

#### Surgery and recording

Mice were anaesthetised with isoflurane and the skull exposed under aseptic conditions. A custom-built stainless-steel metal plate was attached to the skull with dental cement and a metal screw was implanted over the somatosensory cortex on the right hemisphere, for future use as a reference electrode. The skull above SC was left accessible. Animals were allowed to recover from surgery for at least seven days; analgesia was provided for at least the first three days. Mice were then habituated to the experimental apparatus, as described in Visual Stimulus section below. Following habituation, mice were briefly re-anesthetized with isoflurane and a craniotomy was performed over the SC on the right hemisphere, centred at 0.75 mm lateral to sagittal midline, and at lambda), and the dura was left intact. The craniotomy was sealed with silicon elastomer (KwikCast, World Precision Instruments).

Mice were allowed to recover for at least 4 hours before the first recording session. Multiple, daily recordings (3 - 6 sessions, one session per day) were then made in each animal. In each session, the animal was head-restrained in the apparatus, the craniotomy was exposed and a silicon probe, comprising two shanks each with 16 electrodes (spacing 250 µm between shanks, 40 µm between sites; ASSY-37 E-1, Cambridge Neurotech Ltd, Cambridge, UK), was implanted using a vertical micromanipulator (Sensapex, Finland). Electrophysiological signals were acquired using an OpenEphys acquisition board ^51^ at a rate of 30 kHz. The electrodes were lowered rapidly to a depth of approximately 0.5 mm below the brain surface, and then lowered slowly while displaying a large flickering (2 Hz) checkerboard across the visual field. The depth of the dorsal surface of SC was identified by the appearance of robust neural activity modulated at the stimulus frequency; the electrodes were then lowered further until the tip was 500 µm below SC surface (for recordings from SCs) or was 800 µm below (for recordings from SCim). At the end of the recording session the electrodes were retracted and the craniotomy was resealed as above. To confirm electrode targeting the electrode was immersed into a DiI solution before the last recording session. Following that session, mice were deeply anaesthetized and perfused transcardially, and brains were extracted for histological analysis.

#### Visual Stimulus

We used an experimental apparatus described previously ^31^. Mice were head-fixed above a polystyrene wheel (radius 10 cm), such that their head was in the centre of a truncated spherical dome. Visual stimuli were produced by shining the light of a projector (Casio Green Slim XJ-A257-UJ DLP; 60 Hz refresh rate), onto the internal surface of the dome via an hemispherical mirror; the projected image spanned 240° azimuth (from -120° to 120°) and 120° elevation (from -30° and 90°) with mean luminance ∼10 cd.m^2^. Mesh-mapped and gamma-calibrated visual stimuli were produced by the package BonVision ^52^, in the Bonsai framework ^53^. Movement of the polystyrene wheel was sensed with a rotary encoder (2400 pulses/rotation, Kübler, Germany), the output of which was copied to both the OpenEphys acquisition board and to an Arduino connected to the stimulus computer, so as to allow locomotion-dependent updating of the visual scene where required. A synchronising signal was sent to both the OpenEphys board and the stimulus computer. The OpenEphys board also acquired the signals of a photodiode (PDA25K2, Thorlabs Inc., USA) that detected light in a small region of the dome hidden from the animal, and provided additional confirmation of stimulus timing.

We presented three classes of stimulus in the virtual environment: 1) platform: a static virtual platform formed by a smoothed random visual texture (8 cm wide x 100 cm long, rendered 2 cm below the animal’s eye); 2) static object: a round object (8 cm diameter) sitting on the distal end of the platform; 3) looming object: the same round object sitting on the distal end of the platform, but which then moved along the platform towards the animal at one of 12.5, 25, 50 or 100 cm/s, and disappeared after it had collided with the animal. For looms of 100 cm/s, the object appeared on the platform and remained stationary for 1s before moving towards the animal.

Animals were habituated to the virtual environment for 8-13 days (one session per day) before recordings started. On each habituation day the animal was placed in the virtual reality apparatus for up to 30 mins, during which it moved along the virtual platform by running on the treadmill. On each of the last 4 habituation days, the static object was presented at the end of the virtual platform on a subset of randomly interleaved trials. Animals (n = 16) experienced an average 55±30 (mean ± s.d.) trials of the platform, and 52±23 trials with the static object, on each of these days. Each trial was separated by 2s of grey screen. Subsequent recording sessions started with 5 consecutive platform trials, then 40 trials of the static object that were randomly interleaved with 20 trials of a looming object moving at 100 cm/s. In the virtual reality (‘VR’, or ‘closed-loop’) condition, the distance travelled on the polystyrene wheel was used by the stimulus generator to control the position of the animal on the virtual platform. In the ‘replay’ (or ‘open-loop’) condition, the movies produced during the VR condition were replayed to the animal, independent of the animal’s movement on the treadmill. In 2/16 animals, the 40 VR trials and 20 looming object trials were presented in a single block, and these trials then replayed in a single block. In 14/16 animals, we split the 40 trials so that animals experienced 20 trials in VR then 20 trials in replay condition, and then repeated this process. Subsequently, we presented randomly interleaved trials of a looming object moving at different speeds (30 trials/speed); each trial was preceded by 5s of grey screen. To map receptive fields we then obtained responses to flashed black or white squares (15° wide). On each of 3000 0.1s trials (no interstimulus interval), we presented black or white squares at each of 5 random locations drawn randomly from an 8×8 grid centred in the left hemifield. The stimuli covered -15:105° azimuth, and -30:90° elevation, where 0° is directly ahead of the animal and at eye height.

#### Spike sorting and clustering

Electrophysiological signals from all recordings in a session were concatenated and processed using Kilosort 2 and Phy ^54,55^. We kept for further analysis those units in which minimum inter-spike interval was greater than 1 ms, yielding a total of 828 units in SCs (28±10 per session, mean±s.d.; 30 sessions in 7 animals), and 1024 in SCim (29±8 per session; 35 sessions in 9 animals). Subsequent analyses were performed in MATLAB R2021b or 2022a. Neural firing rate was resampled to the refresh-rate of the visual environment (60 Hz). Except for receptive field analyses, these binned rates were transformed into z-scores by normalising to the mean and standard deviation of firing rate across all stimulus conditions. To assess running behaviour, locomotion speed was also resampled at 60Hz and animals were defined as ‘stationary’ if locomotion speed was less than 2 cm/s, and as ‘running’ otherwise.

#### Receptive field analysis

Units were included for further analysis if they produced at least 100 spikes during the stimulus sequence (yielding 711 units in SCs, and 830 in SCim). We analysed responses to black and white stimuli separately. For each unit, a linear regression (function *fitlm* in Matlab) was conducted between a vector representation of the stimulus contrast at each location, and mean firing rate over the frame. The five parameters of a circular two-dimensional Gaussian (x- and y-location, standard deviation, amplitude, and offset) that best predicted the spatial pattern of regression weights were found using constrained least-squares optimisation (function *lsqcurvefit* in Matlab). Size (standard deviation) of the receptive field was constrained to be at least 3.75°. Fits were included for further analysis if the predicted location of the receptive field centre was at least 7.5° from the edge of the stimulus grid, and the fits predicted at least 33% of the variance in the regression weights. Of the 711 SCs units, 569 yielded acceptable fits for black stimuli and 351 for white, of which 38 were acceptable only for white; of the 830 SCim units, 219 yielded acceptable fits for black stimuli and 46 for white, of which 11 were acceptable only for white. Receptive fields could be defined for black squares in most of these units. Since the looming object was also black, we used responses to black squares for further analysis. Note we will overestimate receptive field size, particularly in SCs, because receptive field size was poorly constrained when units responded to only one location in the stimulus grid. The model returned receptive field size less than 5 degrees in 240 SCs units, and 10 SCim units.

#### Response during object loom and object approach

For analyses of responses to object looms in **Fig 1**, binned and z-scored firing rates on each trial were aligned to the time at which the object collided with the animal in the virtual environment. Units were included for further analysis if they met the criteria for response consistency described below.

Response amplitude was calculated as the mean z-score in the 0.2 s (for 100 cm/s objects) or 0.8s (for 25 cm/s) preceding collision. For analyses of responses during object approach in **Fig 2**, the binned and z-scored firing rate was first smoothed with a Gaussian filter (0.3 s). We then discretized the distance between the animal and object into non-overlapping bins of 1cm, and found the time-bins in each trial where the animal was at each of those distances. We then calculated time- and trial-averaged neural activity, and locomotion speed, at each of the 100 distances-to-object. Units were included for further analysis if they met the criteria for response consistency as below.

#### Relative contribution of speed and distance-to-object during object approach

To evaluate the relative contribution of locomotion speed and distance-to-object in explaining SC responses in **Fig 2C**, we used methods previously applied to responses in mouse primary visual cortex ^56^. Distance-to-object was discretised into 110 bins; locomotion speed into 20 bins. We fit a linear model to get weights for each bin of the predictor variables, i.e. *y*_*n*_ = *w*_*n*_*X*, where *y*_*n*_ is the response of the *n*^*th*^ neuron over time, *X* is the matrix of predictor variable bins, and *w*_*n*_ is an array of weights acting on the predictor variable bins to capture the response of the *n*^*th*^ neuron. We used linear ridge regression to calculate the weights based on the equation: 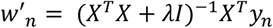, where λ was the ridge parameter tested (one of 0.001, 0.1, 1, 5, 10). We chose λ to minimise cross-validated explained variance. To calculate cross-validated explained variance, we first calculated the weights using the training data (80% of data) and then used the following equation to calculate performance in the test data:

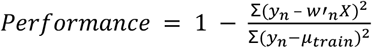

where μ_*train*_ is the mean of the response in the training period. Performance of each of three models (locomotion speed only, distance-to-object only, or both) was calculated on the best λ of each of the models independently.

#### Response consistency

For each of 30 iterations we split trials randomly into two equally-sized sets and binned activity as a function of time to collision (object loom) or distance-to-object (object approach). We averaged activity in each bin over trials and calculated Pearson’s correlation coefficient between the two resultant vectors. We then took one of these sets and randomly shuffled the time- or distance-bins on each trial before creating the average vector, and then calculated the Pearson’s correlation with the other unshuffled, trial-averaged vector as above. The procedure was also repeated 30 times and yielded 30 estimates of raw correlations and 900 estimates of shuffle correlations. Units in which the mean raw correlation exceeded 95% of shuffle correlations were considered consistent.

#### Logistic regression

To classify whether epochs of 0.2 s belonged to a VR or replay condition in **Fig 4**, we performed logistic regression based on purely behavioural covariates or when also including neural population activity in SCim or SCs. We applied control analyses and exclusion criteria to prevent inflated classification performance that could result from temporal structure of the trials or behavioural differences between environments.

##### Inclusion criteria and preprocessing

We included sessions where we ran two repetitions of VR and replay conditions. Running speed strongly influenced neural activity (**Figs 1&2**), and running speed varied between VR and replay conditions in many sessions **(Fig. 3)**. We therefore excluded 5 SCs and 16 SCim sessions during which the animal had very different running profiles or paused for long periods of time during the first replay condition compared to VR conditions (**Suppl. Figure S3**). Missing values in running speed data were imputed using a monotonic cubic interpolation method ^57^. We applied a moving average filter over 81 time steps to correct for slight jitter in wheel movements, followed by a median filter with a kernel of 51 time steps. Neural activity from units that contained missing values were excluded from further analyses.

##### Model details

We divided data into non-overlapping 0.2 s epochs, for each of which we calculated different features (e.g., locomotion speed, z-scored neural activity, etc.) as predictors. We classified these epochs as belonging to VR or replay conditions using logistic regression with sparsity-inducing L1-regularisation (C=1.0). For each session, we trained a separate logistic regression model. We split the data into four train and test splits and report accuracy values averaged over the different data splits (mean and s.d.). We balanced the number of epochs across conditions. We introduced a buffer period between training and test epochs to account for autocorrelations in the neural data and prevent information leakage between training and test sets. Thus, 40 consecutive training epochs were separated from 10 test epochs by a buffer phase of 5 seconds. In control analyses, we tested classifier performance by shuffling test labels between the VR and replay conditions.

##### Feature selection

We first established classification performance for non-neural features (‘behavioural covariates’: locomotion speed, distance to object, time into the experiment), and combinations of these features. Because these covariates could successfully classify many epochs, we applied feature-specific corrections (‘covariates cor.’) to minimise their influence when evaluating the contribution of neural activity to classification accuracy. First, we restricted analyses to epochs where the animal was within 50 cm of the object. Second, we selected epochs to ensure a consistent range of locomotion speed across conditions. Third, to balance average time into the experiment, we only included data from the second half of the first VR condition, the first replay condition, and the first half of the second VR condition (**Suppl. Fig S7**). Note that this control will not be beneficial for more complex non-linear classifiers. Corrected covariates alone still yielded good classification performance in some sessions. We thus concatenated neural features with covariates and assessed the improvement in classification accuracy over that provided by the covariates alone. We performed min-max scaling for each feature (neural and covariate) separately to ensure fair feature weighting.

##### Statistical Testing

To evaluate whether performance accuracy was above chance level for different predictors, we performed a one-sample Wilcoxon signed-rank test comparing the mean accuracy of all sessions to a chance level of 0.5 using the implementation of the scipy.stats package. To assess the difference in prediction performance between models incorporating neural activity and covariates versus covariates alone, we used a paired Wilcoxon signed-rank test. This test compared the mean classification accuracy of all sessions with and without neural activity as a predictor.

##### Code

(Pre)processing, analysis, and classification code was written in Python using the Sklearn toolbox.

## Supplementary Figures

**Suppl. Figure S1:**
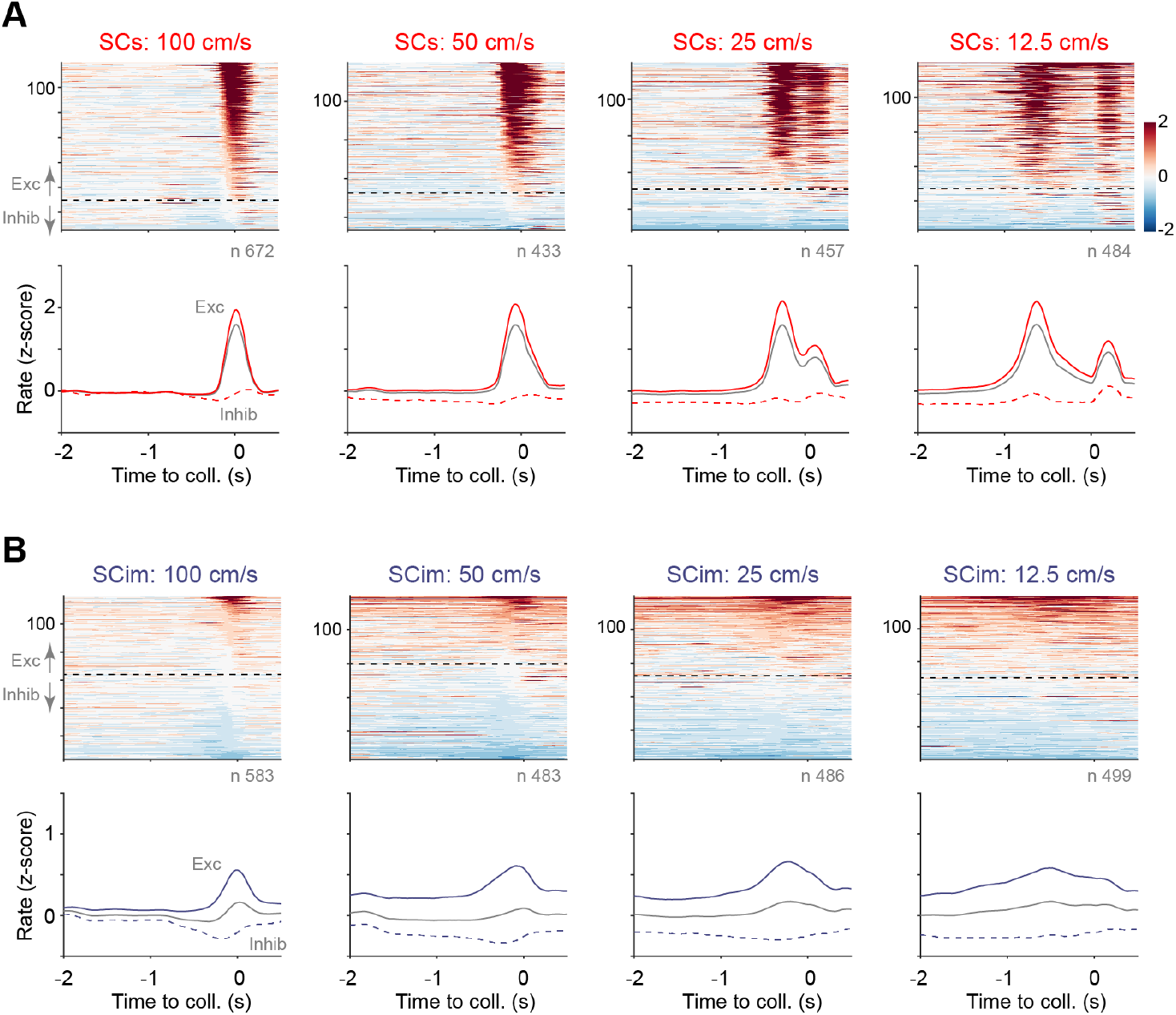
Response of SCs and SCim units to objects looming at different speeds. **A**. Responses of units in SCs to objects looming at the animal at speeds of 100, 50, 25 and 12.5 cm/s (left to right). Conventions as in Figure 1D. At each speed, only units in which the response at that speed was consistent (see Methods) were included. Responses were sorted by the amplitude of the response in the 0.2 s (100 cm/s), 0.4 s (50 cm/s), 0.8 s (25 cm/s), or 1.6 s (12.5 cm/s) period preceding object collision. The second peak, beginning after 0 s at speeds of 25 and 12.5 cm/s, is a visual response triggered by the disappearance of the object after colliding with the animal in the virtual environment. **B**.Same as A but for units in SCim.

**Suppl. Figure S2:**
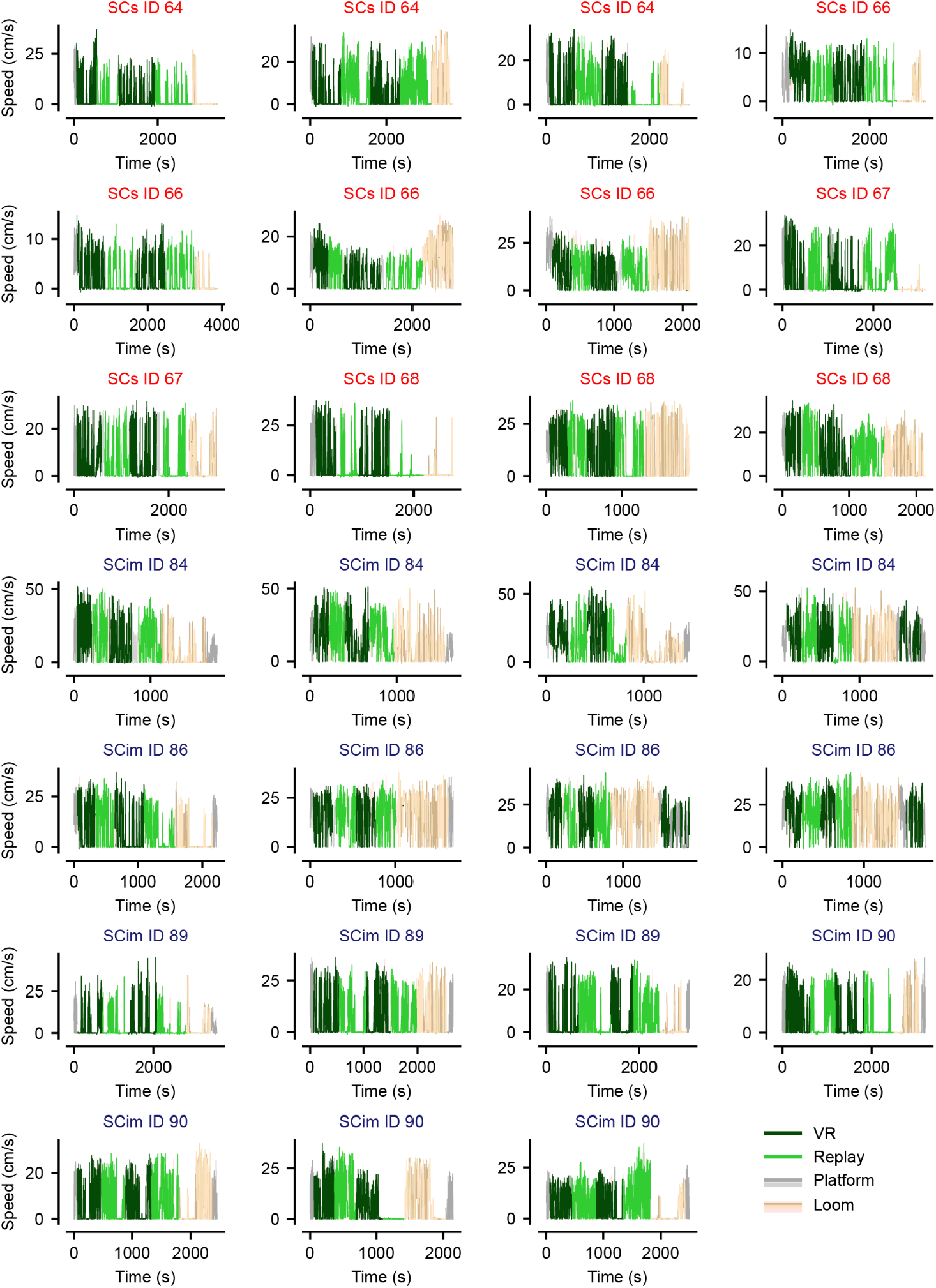
Temporal profile of locomotion speed for sessions where overall locomotion behaviour during Replay condition was similar to that in VR. These sessions were included in the classification analysis. Each plot shows locomotion as a function of time into the recording session. In each case, line colour indicates condition (VR or Replay; Platform; Looming object).

**Suppl. Figure S3:**
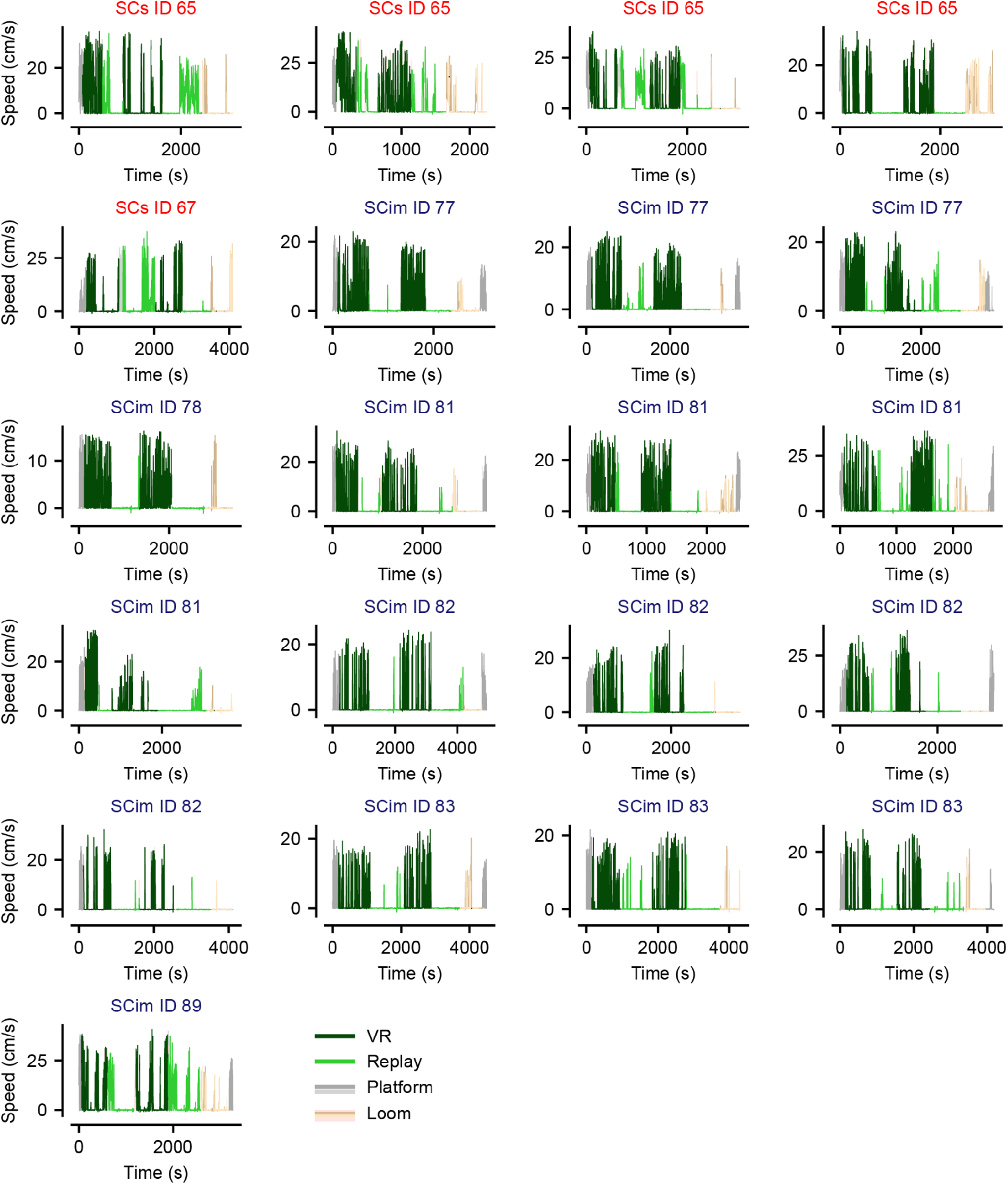
Temporal profile of locomotion speed for sessions where overall locomotion behaviour during Replay condition was very different to that in VR. These sessions were excluded from the classification analysis. Conventions as in Suppl. Figure 2.

**Suppl. Figure S4:**
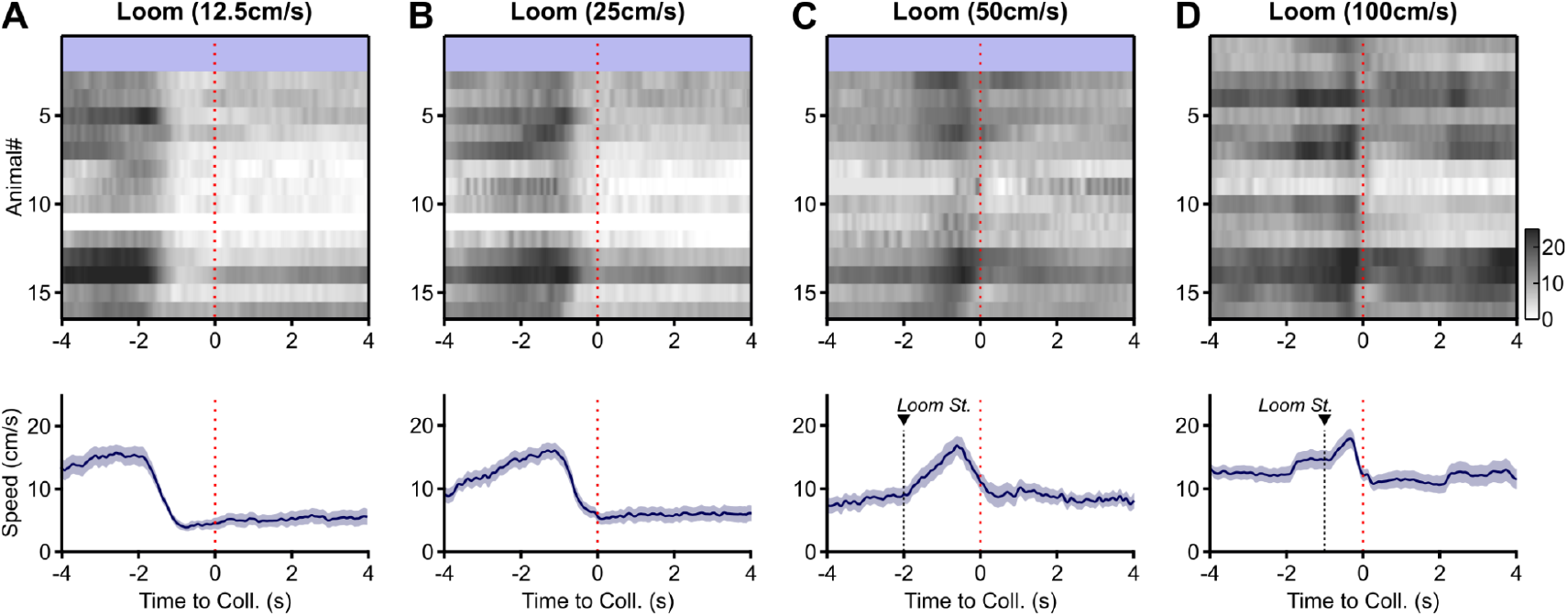
Locomotion speed during presentation of a looming object. **A-D**. Locomotion speeds during presentation of objects looming at 12.5, 25, 50, or 100 cm/s respectively. (*upper*): Locomotion speed for each animal, averaged across trials. Red dashed lines show time of collision. Blue shaded area in **A-C** indicates that the first two animals were not exposed to slower looms. Colorbar in **D** indicates animal speed in cm/s, and applies to all panels. (*lower*): Average locomotion speed across all animals. Red dashed lines show time of collision. Black dashed lines in **C** and **D** (‘Loom St.’) indicate when the object started to move in those trials. For 100 cm/s looms the object appeared and then remained stationary for 1 s before starting to move towards the animal.

**Suppl. Figure S5.**
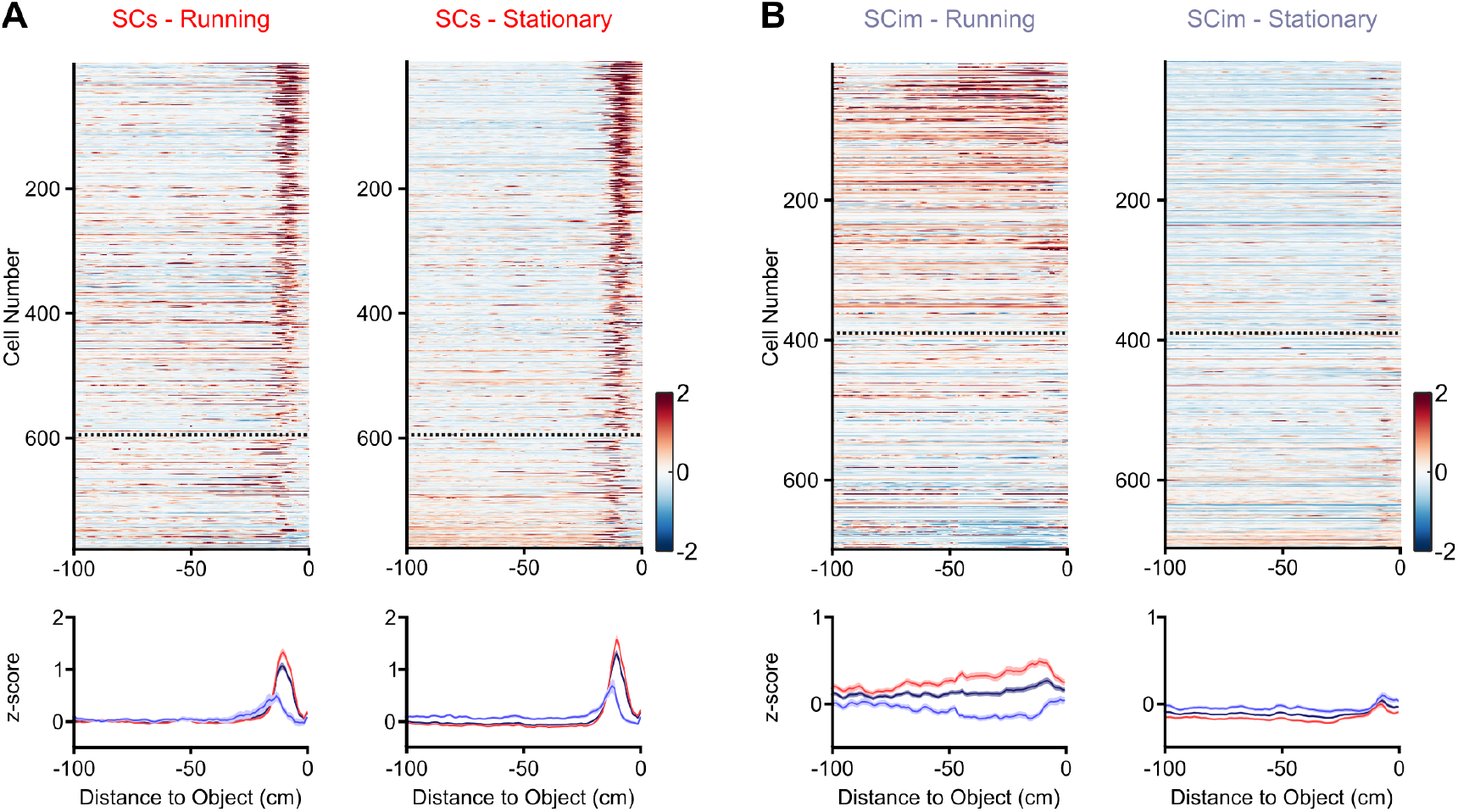
Activity in SCs and SCim during replay of object approach. **A**. Activity of SCs responsive units during replay. Same units as in Figure 2A. Conventions as in Figure 2A. (*upper*) Each replayed trial was classified as presented while the animal was ‘running’ (average locomotion speed >2 cm/s) or ‘stationary’ (locomotion speed <=2 cm/s). Activity was averaged over all running trials (left) or all stationary trials (right). Units are ordered by their response in VR presented in Figure 2A, and the dashed line separating ‘Exc’ and ‘Inhib’ groups is also reproduced from that in VR. (*lower*) Mean ± s.e.m. of activity in running (left) and stationary (right) replay trials, for all units (black), for ‘Exc’ units defined by VR responses (red) and for ‘Inhib’ units defined by VR responses (blue). **B**. Same as A but for units recorded from SCim.

**Suppl. Figure S6:**
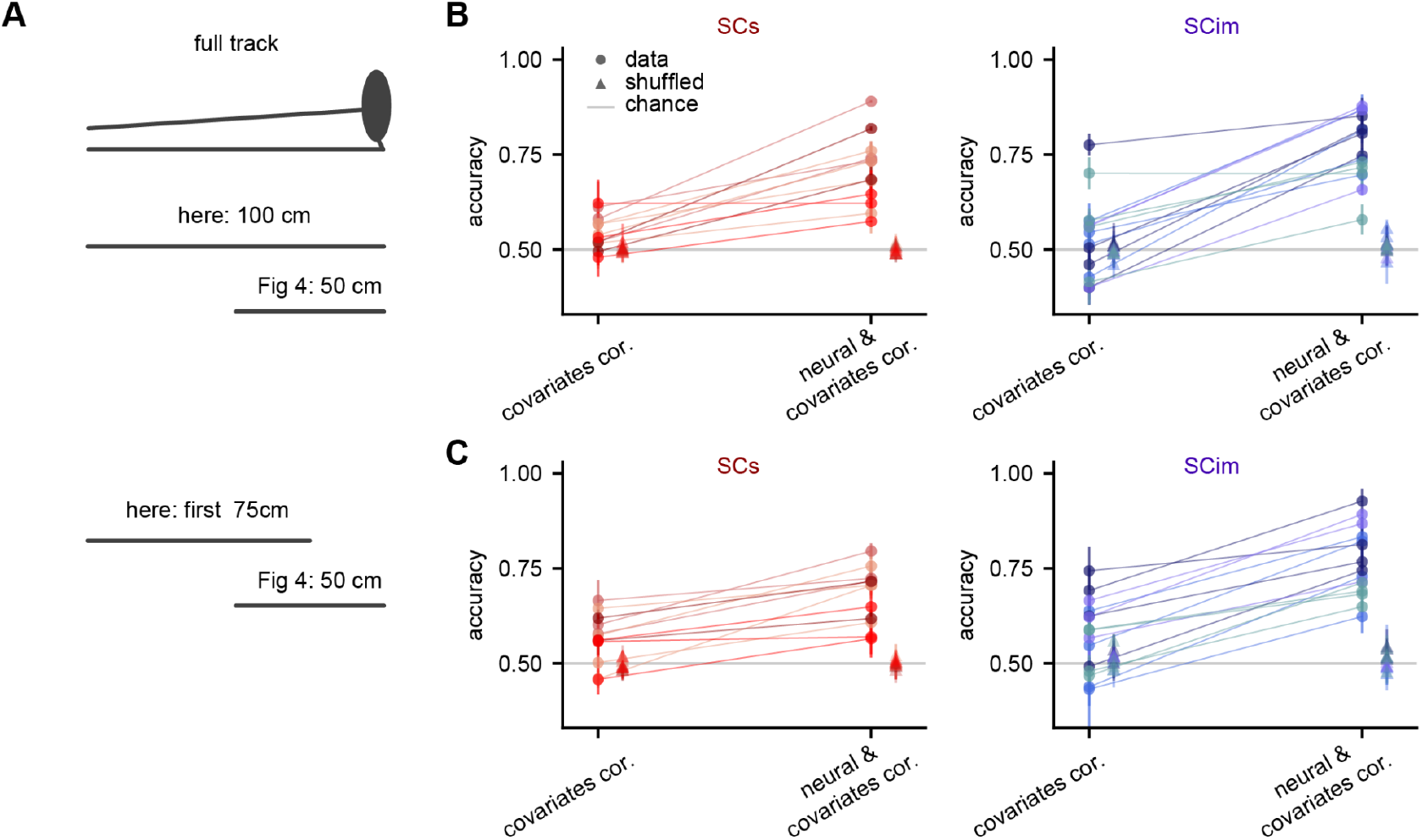
Neural activity also improves classification performance when including data along the entire track. **A**. Schematic of data along the track included in the classification analysis on the right. **B**. Test classification accuracy when considering data of the entire platform (100 cm) of SCs and SCim sessions considering all (corrected) covariates or neural activity and these covariates. The same hue marks sessions from the same animal. Chance level is indicated in grey. **C**. Same as B but for the first 75 cm of the platform, excluding the area closest to the object.

**Suppl. Figure S7:**
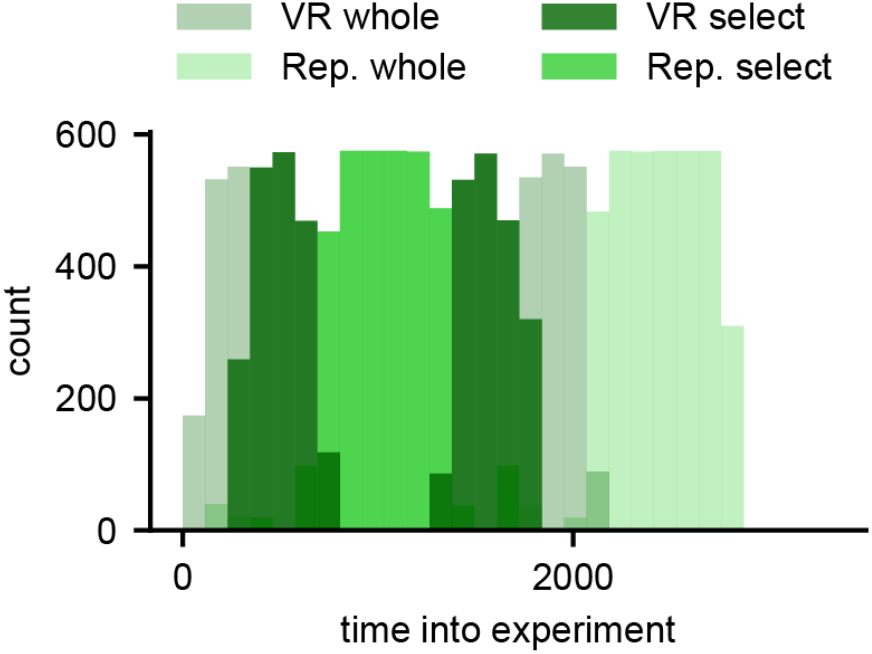
Distribution of time-into-experiment of epochs used in the classification experiment. To ensure that average time-into-experiment was the same in the VR and replay sessions, we selected only parts of the first and second VR condition (‘VR select’) and only the first replay session (‘Rep. select’).

